# Second-generation lung-on-a-chip array with a stretchable biological membrane

**DOI:** 10.1101/608919

**Authors:** Pauline Zamprogno, Simon Wüthrich, Sven Achenbach, Janick D. Stucki, Nina Hobi, Nicole Schneider-Daum, Claus-Michael Lehr, Hanno Huwer, Thomas Geiser, Ralph A. Schmid, Olivier T. Guenat

## Abstract

The complex architecture of the lung parenchyma and the air-blood barrier is difficult to mimic in-vitro. Recently reported lung-on-a-chips used a thin, porous and stretchable PDMS membrane, to mimic the air-blood barrier and the rhythmic breathing motions. However, the nature, the properties and the size of this PDMS membrane differ from the extracellular matrix of the distal airways. Here, we present a second-generation lung-on-a-chip with an array of in vivo-like sized alveoli and a stretchable biological membrane. This nearly absorption free membrane allows mimicking *in vivo* functionality of the lung parenchyma at an unprecedented level. The air-blood barrier is constituted by human primary lung alveolar epithelial cells from several patients and co-cultured with primary lung endothelial cells. Typical markers of lung alveolar epithelial cells could be observed in the model, while barrier properties were preserved for up to three weeks. This advanced lung alveolar model reproduces some key features of the lung alveolar environment in terms of composition, alveolar size, mechanical forces and biological functions, which makes this model a more analogous tool for drug discovery, diseases modeling and precision medicine applications.

---

Organs-on-chips (OOCs) are emerging as predictive tissue modelling tools and as a credible alternative to animal testing. These micro-engineered, cell-based systems provide cells with an environment that closely resembles their native *in vivo* milieu^1,2,3^. Tissue models of physiologically healthy or pathological primary cells from patients have been established, and are robust enough to permit applications such as drug screening^4,5,6^. Micro-engineered systems with an integrated membrane in a microfluidic setting have been reported to model various barrier tissue interfaces, such as those of the lung alveoli, the brain and the gut^7^. By implementing a flexible polymeric membrane in such microfluidic systems, mechanical forces, such as those induced by breathing, could be reproduced^8,9,10^.

An important limitation of these *in vitro* models is the use of an artificial basal membrane made of polydimethylsiloxane (PDMS) seeded with cells for culture. Although PDMS has good elastic, optical and biocompatible properties, it can distort the biochemical microenvironment through high adsorption and absorption levels of small molecules^11^. In addition, the non-biological PDMS differs in important ways from the molecular composition and intrinsic stiffness of the native extracellular matrix (ECM), which is known to affect cellular phenotype and homeostasis^12,13^. A complex ECM environment provides the structural basis for cellular growth, and influences cellular morphology, functionality, differentiation and other traits^14,15^. The role of the ECM in tissue development and function is closely associated with its composition and properties^16^. The replacement of PDMS as culturing membrane with a material made of ECM molecules would therefore be a significant step towards emulating *in vivo*-like tissue barriers and functions.

Hydrogels are currently being used extensively in cell culture systems to recreate the chemical composition and structure of the native extracellular matrix^17,18^. Their intrinsic properties^19^, including mechanical features, chemical composition and porosity, make them ideal candidates to supersede PDMS membranes. However, the creation of thin membranes made of ECM molecules, with stretchable properties to mimic the cyclic mechanical strain of the lung alveolar barrier, is technically challenging and has not yet been achieved. Lo and colleagues reported a thin, cellularised collagen membrane integrated into a microfluidic-based blood oxygenator being developed as an extracorporeal lung support^20^. More recently, collagen membranes have been integrated into microfluidic devices for use as cell culture substrates. These membranes were either cast^21,22^, compressed^23^ or spin-coated^24^ on a PDMS surface prior to being sandwiched between two microfluidic structures. The resulting thicknesses of these membranes, not designed to be stretched, were between 15 and 30μm. Harris and colleagues reported the use of a stretchable collagen membrane to determine the mechanical properties of a monolayer of cells. They found that the mechanical properties of the layer were dependent on the integrity of the actin cytoskeleton, myosin and intercellular adhesions interfacing adjacent cells^25^. Dunphy *et al.* added elastin to collagen and developed a stretchable and soft membrane for tissue engineering. However, with a thickness of about 1mm, it was developed to evaluate the mechanical properties of the material and not to mimic the air-blood barrier^26^.

Here, we report a unique, biological, thin and stretchable air-blood barrier made of collagen and elastin. Unlike all other lung alveolar models reported so far, an array of tiny stretchable alveoli with physiological dimensions is reproduced. A thin gold mesh with a pore size of 260µm is used as the scaffold, supporting the array of 40 alveoli. The uncomplicated production process of the membrane allows straightforward modifications of the system that permit a wide variety of investigations of physiological and pathological phenomena. The membrane is created by drop-casting a collagen-elastin (CE) solution onto the gold mesh, where it spreads and is maintained by surface tension (Fig. 1). The resulting membrane is stable and can be cultured on both sides for weeks. Its permeability further allows cells to be cultured at the air-liquid interface, and its elastic properties mimic the respiratory motions by mechanically stretching the cells. Results with primary human alveolar epithelial cells from patients co-cultured with primary human lung endothelial cells demonstrate that the air-blood barrier functions can be maintained and used experimentally in a resilient and reproducible manner. This proto-physiological membrane opens the way to a new generation of lung-on-a-chip and OOC devices that enable the mimicry of biological barriers with a new level of analogy to whole organ systems.

**Figure 1:**
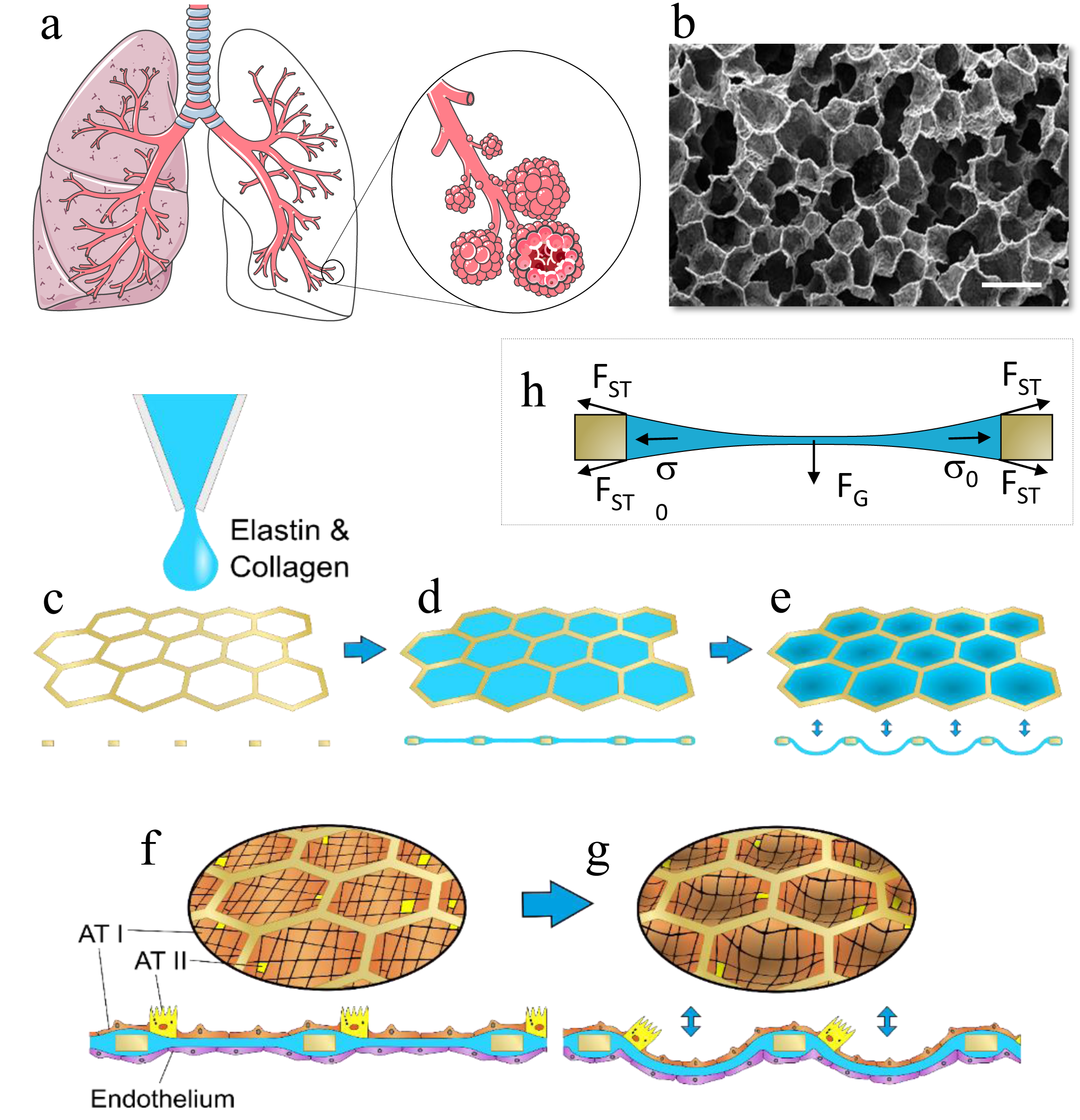
(a). Schematic of the respiratory tree-like structure ending with alveolar sacs (adapted from http://smart.servier.com/). (b). SEM picture of a slice of human lung parenchyma with tiny lung alveoli and their ultra-thin air-blood barrier (courtesy of Prof. Dr. Peter Gehr, Institute of Anatomy, University of Bern; scale bar: 500µm). (c-d). Schematic of the production of the CE-membrane used in the 2^nd^ generation lung-on-a chip. A thin gold mesh with an array of hexagonal pores of about 260µm is used as a scaffold, on which a drop of collagen-elastin solution is pipetted. (e-g). The collagen-elastin gel forms a suspended thin membrane that can be stretched at the alveolar level by applying a negative pressure on the basolateral side of the membrane. (f-g). Type I (ATI) and type II (ATII) primary human lung alveolar epithelial cells are co-cultured with lung endothelial cells on the thin collagen-elastin membrane. (h). Schematic of the force balance during the drying of the membrane. F_ST_, F_G_ and σ_o_ stand for surface tension force, gravity and residual stress, respectively.

## Results

### Production of a thin, biological and stretchable membrane

A simple process was used to create the thin biological membrane (Fig. 1). A drop of CE solution was pipetted onto a 2mm-diameter and 15µm-thin gold mesh (Fig. 1C and 2A) made of an array of 40 regular hexagons, with sides of 130µm separated by 30µm-wide walls. Once pipetted onto the mesh, the CE drop was maintained on its top by surface tension forces. After a gelation step at 37°C, the CE solution dries out at room temperature within two days. While water evaporates from the drop, surface tension forces and residual forces counteract gravity force enabling the suspended membrane to form (Fig. 1H). Figures 2B-D illustrate the dried CE-membrane with a thickness of only a few micrometers that is suspended on the hexagons array. Once dried, the membrane was integrated into a microfluidic chip, where it was sandwiched between two microfluidic parts, a top part in PDMS with an apical reservoir and a bottom part in polycarbonate that formed the basolateral chamber (Sup. Fig. 1). The dried membranes are robust and can be stored for at least 3 weeks at room temperature. The membranes are rehydrated by submersion in cell culture medium for 2h prior to cell seeding.

**Figure 2:**
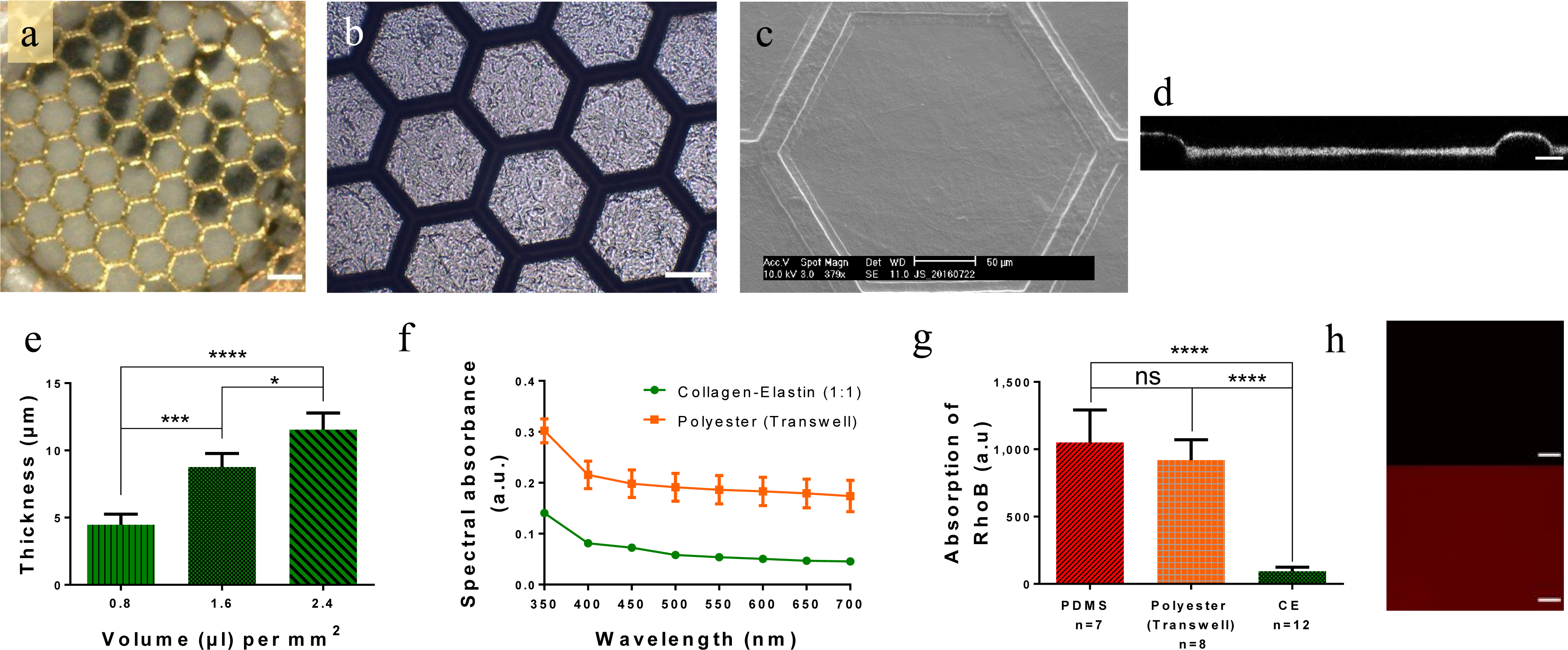
Properties of the thin biological membrane. (a). Optical clarity of a 10µm-thin CE-membrane integrated in the gold mesh. Scale bar: 200µm. (b). Picture of an array of several hexagons with a CE-membrane. Scale bar: 100µm (c). SEM picture of the CE-membrane suspended in one hexagon. (d). Cross-section of the CE-membrane visualized via confocal microscopy. Scale bar: 20µm. (e). Characterization of the CE-membrane thickness in function of the collagen-elastin solution volume pipetted on top of the gold mesh. (f). Comparison of the spectral absorbance of the CE-membrane and of a polyester membrane (Transwell insert 0,4µm pores sizes). (g). Difference of Rhodamine B (10µM) absorption between a 10µm-thin CE-membrane, a 10µm-thin PDMS membrane and a polyester porous membrane (Transwell insert, 0.4µm pores sizes). (h). Pictures of CE-membrane (top) and a PDMS membrane (bottom) after being exposed to RhoB for two hours. Scale bar: 200µm.

### Properties of the CE-membrane

The thickness of the membrane was evaluated using reflective light. With a CE ratio of 1:1, the thinnest membrane obtained had a thickness of 4.5 ± 0.8µm for a pipetted CE solution volume of 0.8µL/mm^2^ (Fig. 2E). When the pipetted volume was doubled (1.6µL/mm^2^), the thickness of the membrane also doubled (8.8 ± 1µm). A thickness of 11.5 ± 1.2µm was obtained with 2.4µL/mm^2^. Decreasing the elastin concentration (2:1 ratio) resulted in a reduction of the membrane thickness (Sup. Fig. 2), to the detriment of its viscoelastic properties (Fig. 3C). The membrane thickness was homogeneous within each hexagon. Variation in membrane thickness across the array was less than 20% (Sup. Fig. 3) with a pipetted volume of 1.6µL/mm^2^. Confocal (Fig. 2D) and transmission electron microscopy (TEM) imaging (Fig. 7B) of the membrane cross-section confirmed these findings.

**Figure 3:**
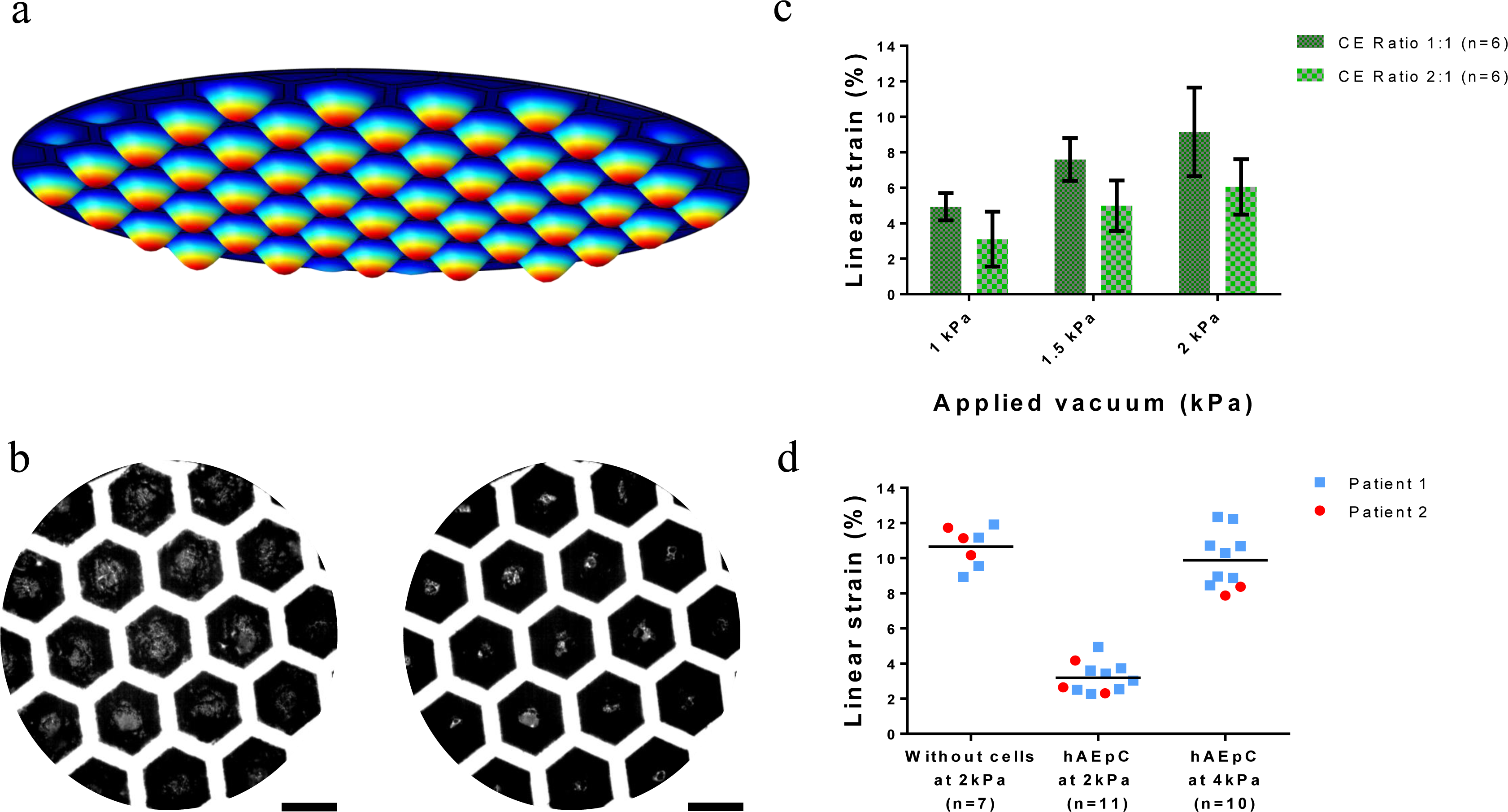
Membrane flexibility. (a). Numerical simulation of the deflection of the CE-membrane array. (b). Picture of a CE-membrane array at rest (left) and exposed to a negative pressure of −2kPa (right). Scale bar: 200µm. (c). Linear strain inside the small alveoli in function of the applied vacuum and the composition of the CE-membrane. (d) Maximum linear strain without (control) and with confluent lung alveolar epithelial cells of two patients at two pressures.

The optical properties of the CE-membrane were assessed by light spectrometry. The CE-membrane performed better than a polyester (PET) membrane of standard Transwell inserts. The 10µm-thin CE-membrane absorbed about 10% of visible light, whereas a 10µm-thin PET membrane with 0.4µm pores used in inserts absorbed about 20% (Fig. 2F). This low absorbance level was also obtained for a 2:1 ratio CE-membrane and for a collagen membrane (Sup. Fig. 4). The excellent optical properties of the CE-membrane were qualitatively confirmed by text placed at the backside of the membrane that was easily read from the apical side (Fig. 2A).

Absorption and adsorption of small molecules on the membrane were tested using exposure to rhodamine B. Compared with PDMS and with the PET membranes of similar thicknesses, the CE-membrane absorbed much less rhodamine B. After 2h of immersion in 10µM rhodamine B, the number of fluorescent molecules ab/adsorbed was about 90% lower in a 10µm-thin CE-membrane than in the PDMS and the PET membranes (p-value < 0.0001) (Fig. 2G). The absorptions/adsorptions of all polymeric membranes tested are higher than those of all biological membranes (Sup. Fig. 5). This absorption difference is illustrated in Figure 2H, which shows the PDMS and CE-membranes after 2h incubation time with rhodamine B. Using the same imaging setting parameters, the PDMS membrane absorbs more rhodamine B than the CE-membrane.

The stretchability of the CE-membrane was tested by applying a cyclic negative pressure to the basolateral chamber. The membranes of the 40 hexagons deflect simultaneously and homogeneously in three dimensions (Fig. 3B and Sup. Fig. 6). For the 1:1 CE-membrane, the applied radial strain reaches 4.9% ± 0.8% for a negative pressure of 1.0kPa, and almost doubles (9.2% ± 2.5%) when −2.0kPa is applied (Fig. 3C). Figure 3A shows a numerical simulation of the deflection of the membranes in the array of hexagons. When the elastin concentration was decreased, the membrane became stiffer, which resulted in smaller linear strains. For example, at −1.5kPa, the radial strain was 5.0% ± 1.4% for a 2:1 ratio, whereas it attained 7.6% ± 1.2% for a 1:1 ratio (Fig. 3C). The gold mesh slightly deflected during the experiments, but this did not influence the individual deflection of the membrane in each hexagon (Sup. Movie). When lung alveolar epithelial cells were seeded onto the membrane, 4.0kPa was needed to induce a 10% linear mechanical strain (Fig. 3D).

The CE-membrane permeability was assessed by the apical-basolateral transport of two molecules with different molecular weights: FITC-Sodium (0.4kDa) and RITC-Dextran (70kDa). After 4h of incubation, 25.5% ± 4% of the smaller molecules and 12.0% ± 3.7% of the larger molecules were detected in the basolateral chamber (Sup. Fig. 7). The permeability of the membrane was further tested by culturing cells at the apical side of the membrane at the air-liquid interface. In these culture conditions, the nutrients diffuse from the basolateral to the apical side of the membrane. Lung alveolar epithelial cells were successfully cultured at the air-liquid interface for several days (Fig. 4 and Sup. Fig. 8). The cells were confluent and created a functional barrier (see below).

**Figure 4:**
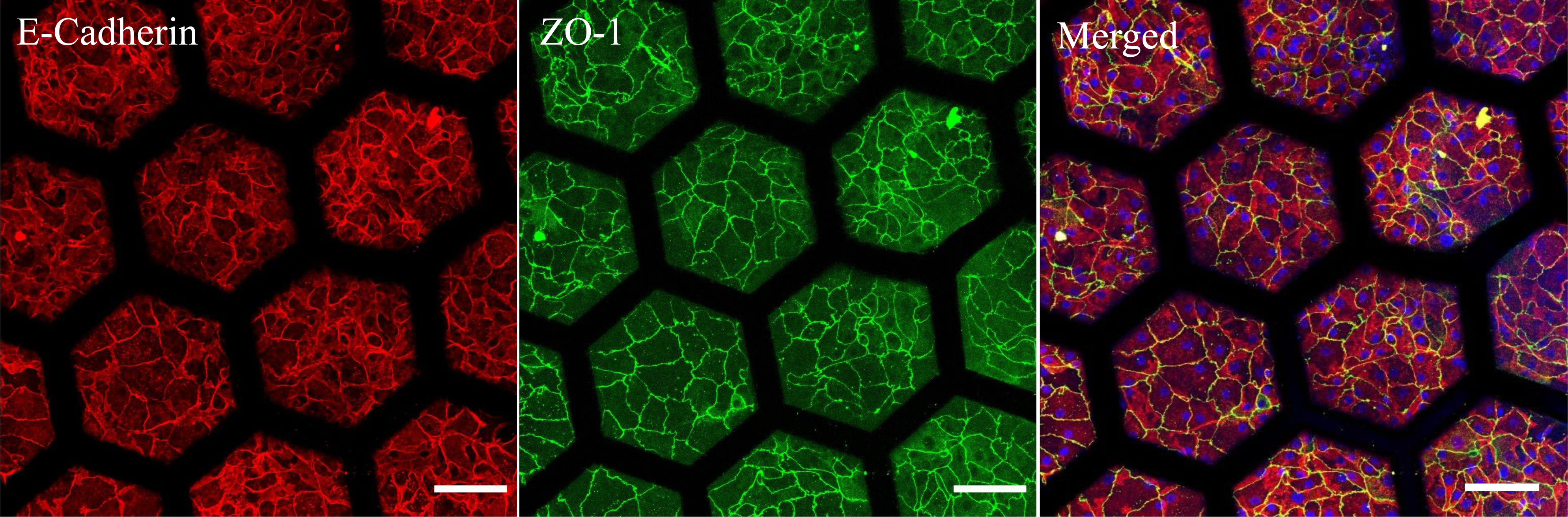
Immunostaining of primary human lung alveolar epithelial cells. hAEpC cultured on the hexagonal mesh with the CE-membrane after 4 days and at air-liquid interface for 2 days with expression of adherent junction markers (E-Cadherin), tight junctions with zonula-occludens-1 (ZO-1) and merged. Scale bar: 100µm.

### The CE-membrane, a good cell culture support

Human primary alveolar epithelial cells (hAEpCs) and human lung microvascular endothelial cells (VeraVec) were successfully cultured on each side of the membrane. The cells tightly adhered to the collagen and the elastin as illustrated in the TEM picture of the membrane cross-section (Sup. Fig. 9). Lung epithelial cells seeded at high and low density spread and proliferated on the membrane (Fig. 5A and Sup. Fig. 10). A significant difference in cellular surface area was observed between day 2 and day 8 between high and low seeding concentration (p < 0.01) (Fig. 5B). At high seeding density, cell confluence was reached at day 2. The cellular surface area remained at 1400 ± 160µm^2^, whereas it increased to almost 2500 ± 136µm^2^ at low seeding concentration. After 2 weeks, primary human lung alveolar epithelial cells were confluent showing nice cell-cell contacts and microvilli (Fig. 5E, F). Primary human lung alveolar epithelial cells and primary human endothelial cells could both be cultured for at least 3 weeks (Sup. Fig. 11).

**Figure 5:**
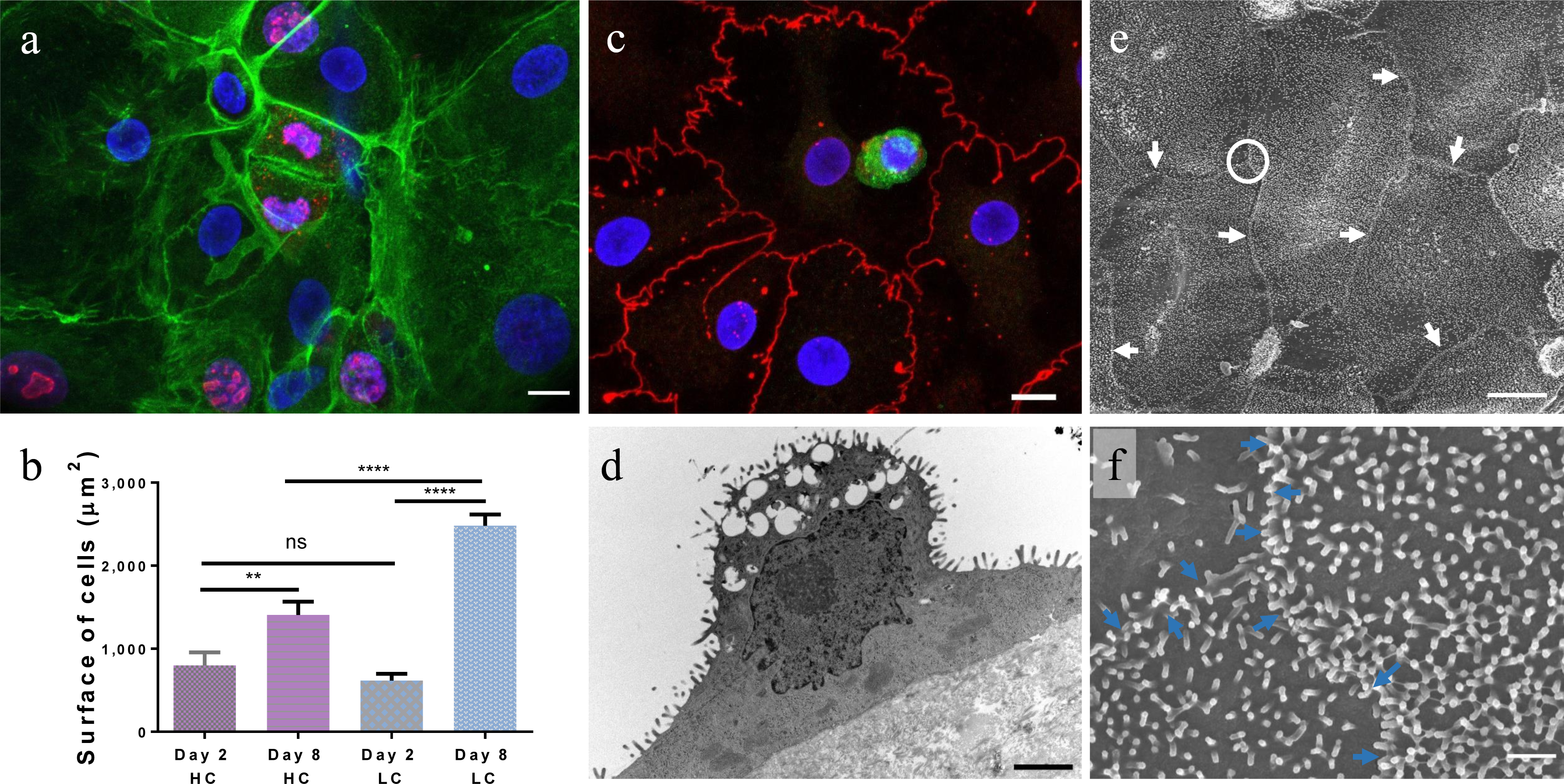
Primary lung alveolar epithelial cells. (a). Expression of Ki-67 marker on hAEpC at day 4. Actin (green), Ki-67 (red) and Hoechst (blue). Scale bar: 10µm. (b). Cellular surface of hAEpC at day 2 and 8 in function of the cell seeding concentration. LC: low seeding concentration (100’000 cells/cm^2^) and HC: high seeding concentration (270’000 cell/cm^2^). (c). Expression of surfactant protein-C (SP-C, green), tight junction (Z0-1, red) and nuclei (Hoechst, blue) at day 4. Scale bar: 10µm. (d). TEM picture of a hAEpC type-II-like cell at day 4, showing its microvilli and empty spaces, where lamellar bodies were located. Scale bar: 2µm. (e) SEM picture of hAEpC at day 14, illustrating tight cell-cell contacts. White arrows: cells border; white circle: area zoomed in (f). Scale bar: 10µm. (f) Intersection between three cells at day 14, showing their interface and a multitude of microvilli. Blue arrows: cells border. Scale bar: 1µm.

### Reproduction of the lung alveolar barrier

The typical phenotypes of lung alveolar epithelial cells were investigated using TEM imaging and immunostaining. The characteristic morphologies of type I (ATI) and type II (ATII) lung alveolar epithelial cells – flat and elongated for ATI (Fig. 7B), small and cuboidal for ATII^27^ (Fig. 5D) – were recognisable by TEM imaging. Tight junctions, a further characteristic of lung alveolar epithelial cells, were clearly identifiable in Figure 6C. Zonula occludens (ZO-1) were expressed along the cell borders at day 4 (Fig. 5C) and day 21 (Sup. Fig. 11). Surfactant protein-C (SP-C) and lamellar bodies, both typical ATII markers, are shown in Figure 5C, and Figure 5D, respectively.

**Figure 6:**
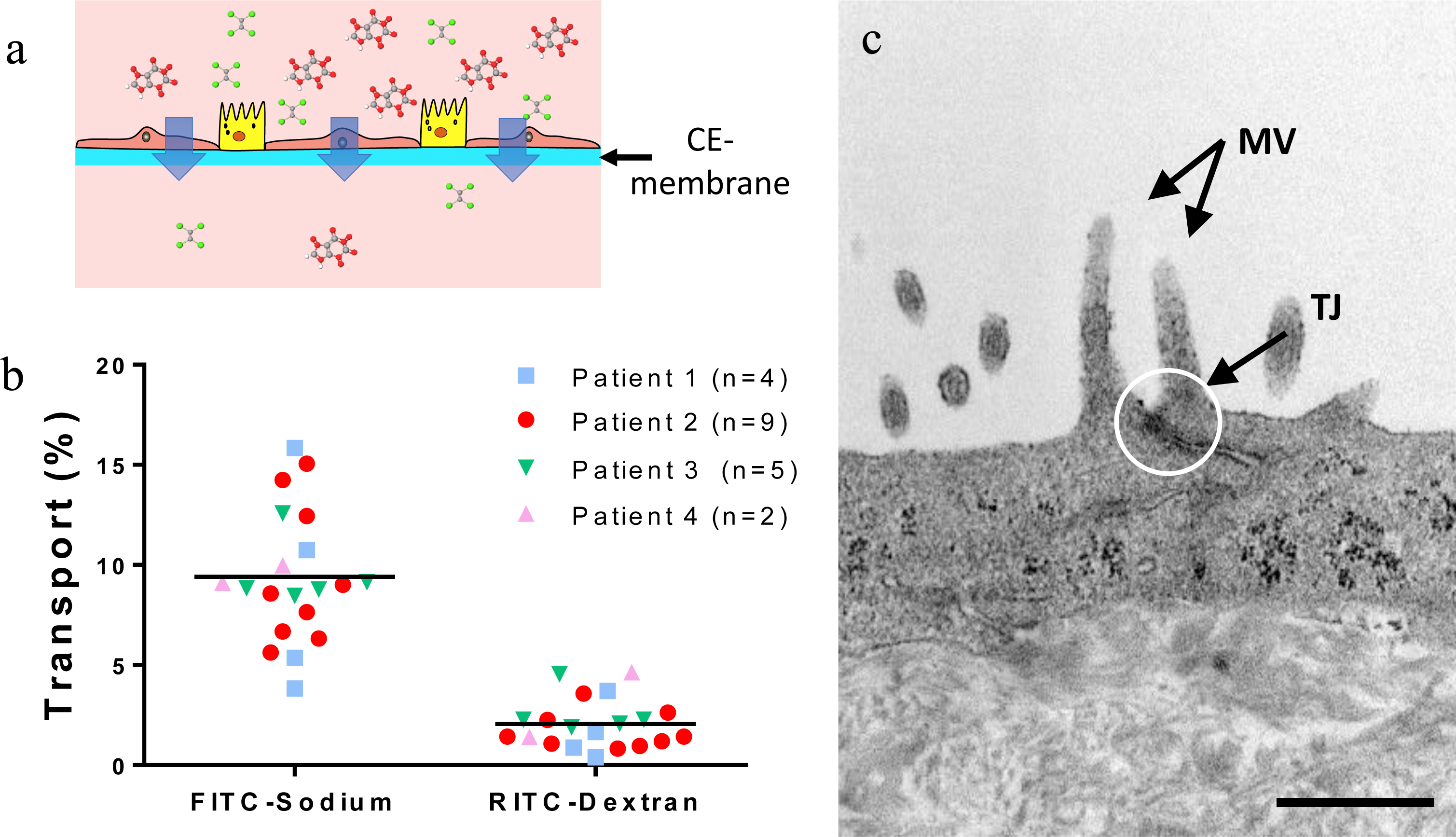
Barrier function. (a). Schematic of the transport of molecules across the CE-membrane cultured with alveolar epithelial cells. (b) Transport of FITC-Sodium and RITC-Dextran molecules across the CE-membrane after 4h of incubation with hAEpC. The experiments were carried with cells from four patients. (c). TEM picture of tight junction (TJ) between two hAEpC. Apical microvillis (MV) typical to type II alveolar epithelial cells can clearly be seen. Scale bar: 500nm.

**Figure 7:**
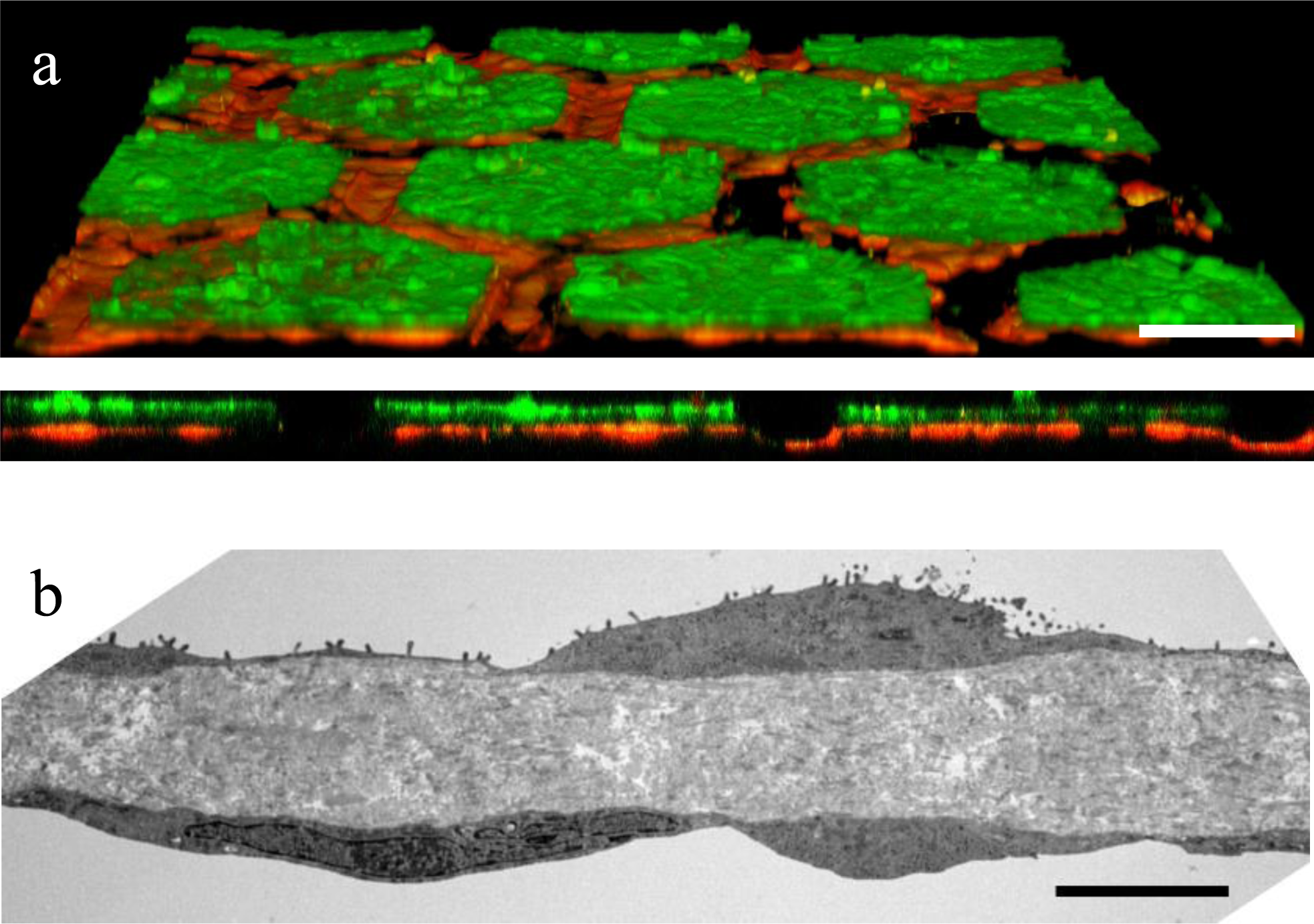
Air-blood barrier reproduction. (a). Confocal pictures (perspective view and cross-section) of a co-culture of hAEpC (E-Cadherin, green) with human primary endothelial cells (Rfp-label, red) on the hexagonal mesh with the CE-membrane. Scale bar: 100µm. (b). TEM picture of hAEpC type-I-like cells in co-culture with human lung endothelial cells at day 4. Scale bar: 5µm.

The permeability of the CE-membrane with a monolayer of lung alveolar epithelial cells was further assessed by testing the diffusion capacity of the two molecules used earlier (FITC-Sodium, RITC-Dextran) as show in Figure 6A. The experiment was performed between days 5 and 8 to guarantee the confluence of the epithelial layer. The transport properties of the membrane were significantly affected by the presence or absence of cells. For FITC-Sodium and RITC-Dextran molecules, 24.7% ± 3.5% and 12.3% ± 2.9%, respectively, were transported through the membrane without cells against 9.4% ± 3.2% and 2.1% ± 1.2%, respectively, with hAEpCs. This result was confirmed with cells from four patients (Fig. 6B).

To further reproduce the lung alveolar barrier, human lung microvascular endothelial cells were cocultured on the basolateral side of the membrane, with lung alveolar epithelial cells on the apical side. Both cell types reached confluence and populated the whole array (Fig. 7A). Figure 7B illustrates a close-up of the alveolar barrier, with the CE-membrane sandwiched between the alveolar epithelium and the microvascular endothelium.

## Discussion

The lung parenchyma comprises of a large number of tiny alveoli organised in a three-dimensional architecture. Thin alveolar walls made of capillary networks and connective tissue separate the alveoli and stabilise the parenchymal construction^27,28^. This complex and dynamic environment makes the lung alveolar unique and difficult to mimic *in vitro*. First-generation lung-on-a-chip devices imitate the rhythmic mechanical strain of the alveolar barrier induced by breathing motions^8,9^. Although these systems allow investigation of the mechanobiology of the air-blood barrier for the first time, they are limited by the nature of the PDMS membrane they are made of. The main drawback of PDMS is that it is synthetic which limits its function and the ability to mimic physiological capacities. The ECM of the lung alveolar region has structural and mechanical cell substrate functions but beyond that the ECM is pivotal in determining normal cellular function and differentiation in health and dysregulation in disease^12,29,30^. Another limitation of PDMS membranes is the absorption and adsorption of small molecules and the effect on the ECM as local reservoir of growth factors and bioactive molecules, which are not maintained by the microenvironment at physiological concentrations, and therefore distort effects in the system. This is also a major concern for preclinical drug testing applications, as the effective drug concentration that cells are exposed to is difficult to evaluate^11^. A further drawback is the rather laborious and challenging fabrication process of ultrathin and porous PDMS membranes^9,31^. In addition, first-generation lung-on-a-chip devices imperfectly reproduce the geometric dimensions of the native lung alveoli, as the surface of the culturing membrane creates a unique alveolus of non-physiological dimensions, rather than an array of alveoli of *in vivo*-like anatomy. This limits investigations of structural and biomechanical changes of alveoli such as those observed in the formation of emphysema^29^.

Here, we present a second-generation lung-on-a-chip with an array of alveoli and a stretchable biological membrane that mimics *in vivo* functionality at an unprecedented level. The CE-membrane reproduces the composition and geometrical, biophysical, mechanical and transport properties of the lung alveolar barrier^28^. It recreates the native viscoelastic microenvironment of the cells. Collagen I, the most abundant type of collagen present in connective tissue^32^, provides structural stability for the alveoli, and elastin adds elasticity, which is essential for withstanding continuous breathing motions. By tuning the CE ratio and/or adding other ECM molecules, scaffolds stiffness can be tailored to specific applications^33^, which is required to model healthy and diseased alveolar environments, such as those present in lung fibrosis^34^. TEM pictures reveal remarkable adhesion of the cells to the CE-membrane and the reproduction of the epithelial/endothelial barrier. The membrane enables the diffusion of small and larger molecules (FITC-Sodium and RITC-Dextran) and of epithelial cell nutrients necessary to culture cells at the air-liquid interface, their physiological microenvironment. The results obtained using cells from four patients were similar. Importantly, the absorption and adsorption issue observed with the PDMS membranes is almost absent.

The hexagonal gold mesh with a suspended CE-membrane provides cells with small alcoves containing an environment similar to that found in an alveolus as measured by a number of different parameters. First, the size of each small alcove is the same order of magnitude as the diameter of lung alveoli, reported to be around 160–200µm^35,36^. Second, the borders of the alcoves mimic the alveolar walls^37,38^ that separate alveoli from each other and strengthen the stability of the structure. Third, the three-dimensional mechanical stress created within each alcove is distributed in a physiological strain gradient. This environment, combined with the CE-membrane, gives the cells more physiological cell culture conditions and may also enable the recreation of biological events that at their onset only involve a limited number of cells. For example repetitive microinjuries of the epithelium that are believed to trigger idiopathic lung fibrosis are a low-cell number phenomenon that could be mimicked^39^. Investigations of phenotypic changes underlying lung cell pathologies and their effect on downstream signalling cascades become possible in tissue-specific primary cell culture microenvironments.

The simple and reproducible production process of the CE-membrane makes it an easy to use tool for academic laboratories as well as for larger scale applications, like screening. The unique gold mesh also allows the creation of larger cell culture surfaces for specific read-outs requiring larger number of cells. The CE-membrane has great versatility as thickness can easily be tuned by adapting the volume of the CE solution pipetted onto the gold mesh to suit any number of experimental requirements. The thinnest membrane obtained has a thickness comparable to the thinnest porous PDMS membrane reported thus far^10^. Unlike synthetic polymers, such as PDMS, the optically transparent CE-membrane does not require any preliminary coating prior to cell seeding. Moreover, the dehydrated extracellular matrix array is robust and can be stored for several weeks at room temperature. Ease of use is further improved as the membrane is mainly created by surface tension force and does not require clean room conditions. Taken together, these characteristics make the CE-membrane a credible alternative to PDMS, with advantages of usability, production and stability.

We have developed a lung alveoli array that displays characteristics of the lung parenchyma with analogy to native alveolar tissue in a number of physiological parameters. Three key features of production and properties were considered in the development of this new generation of organ-on-a-chip. First, a suspended culturing membrane was created by surface tension force. Second, the CE membrane mimics the native ECM of the lung parenchyma. Third, an array of alveoli with more physiological geometric proportions was created by the gold mesh. Replacing less than optimal PDMS membranes is desirable in *in vitro* barrier models, and this makes our CE membrane a versatile and generic solution that can be expanded to mimic other barrier structures found *in vivo*. The robustness and absorption-free membrane properties make the CE-membrane a potentially powerful tool for drug testing, lung diseases modelling and precision medicine applications.

## Materials and Methods

### Production of the CE-membrane

The CE-membrane was produced as follows (Fig. 1). The membrane was based on rat-tail collagen type I, high concentration (Corning, New-York, NY, USA), and bovine neck elastin powder/lyophilised (Sigma-Aldrich, Buchs, Switzerland). The two molecules were mixed at a final concentration of 3.5 mg.mL^-1^ in a pH 7.4 buffer. A 15 µm-thin gold mesh (Plano GmbH, Wetzlar, Germany) with hexagonal pores of 225 μm (inner diameter) and 260 μm (outer diameter) was used as a scaffolding to create the biological membrane. The gold mesh was successively treated with 5% 3-aminopropyl triethoxysilane (APTES) (Sigma-Aldrich) and 0.1% glutaraldehyde (Sigma-Aldrich) to ensure attachment of the membrane. The CE solution was pipetted directly on top of the gold mesh. Its thickness was tuned by adapting the volume of the CE solution pipetted. After pipetting, the chip was immediately placed at 37°C, 100% humidity and 5% CO_2_ for 2 h 15 min to allow gelation of the membrane. Then, the membrane was placed for 48h at room temperature to dry. Membranes were stored at room temperature. Before use, membranes were rehydrated with cell culture media for 2h at 37°C.

### Micro-device fabrication

To create the air-blood barrier-on-a-chip, a polydimethylsiloxane (PDMS Silgard 184, Dow Corning, Midland, MI, USA) plate was attached to a polycarbonate bottom with double tape (Arcare 90445-5, Adhesives Research, Glen Mark, PA, USA). The gold mesh with the CE-membrane was sandwiched between the two chambers (Sup. Fig. 1). This design enabled the compartmentalisation of the culture medium in the apical and the basolateral chamber. The top layer was produced by PDMS soft lithography. Briefly, a prepolymer was mixed with a curing agent at a weight ratio of 10:1 and placed in a vacuum chamber to remove air bubbles. After degassing, the PDMS mixture was cast in a mould with two dowel pins located at the border of the chip as alignment features. After an incubation at 60°C overnight, the PDMS mixture was fully cured, and cut into a rectangular shape of 20 × 15 × 3.2 mm. The bottom layer was made of polycarbonate with a central hole of 2mm and two 1.5mm additional holes on both sides of the lower part to allow access to the membrane. The top layer can easily be detached from the bottom layer to reduce the focal distance required for confocal imaging. Prior to being used, the chip was sterilised by autoclaving (CoolCLAVE, Genlantis, San Diego, CA, USA).

### Cell culture

Primary hAEpCs were isolated from patient tissue according to a protocol reported previously^10,40^. Briefly, alveolar epithelial type II (AT II) cells were isolated from tissue obtained from healthy areas removed from patients undergoing lung tumor resection surgery. All patients gave informed written consent for usage of surgical material for research purposes, which was approved by ethical committee from the Ärztekammer des Saarlandes. All procedures were carried out in accordance with institutional guidelines from Saarland (Germany) and from the Canton of Bern (Switzerland). Cells were cultured in Small Airway Growth Medium (SAGM(tm), Lonza, Basel, Switzerland) with BulletKit (CC-3118, Lonza), supplemented with 1% FBS (Sigma) and 1% P/S. RFP-labelled human lung microvascular endothelial cells (VeraVec, Angiocrine Biosciences Inc., San Diego, CA, USA) were cultured in EGM2 medium (Lonza) supplemented with growth factors according to the manufacturer’s instructions (EGM2-MV BulletKit, Lonza). All cell manipulations were performed in a sterile flow hood, and cells were maintained at 37°C, 100% humidity and 5% CO_2_.

For monoculture, hAEpCs were seeded with a density of 270,000 cells/cm^2^ or at 100,000 cells/cm^2^ (low concentration condition). The cells were incubated for 24h, allowing the cells to adhere to the membrane, and reached confluence after 48h. To create a coculture, the chip was flipped, and VeraVec cells were seeded on the basal side of the CE-membrane at 1.0e^6^ cells/mL. After 24h, the chip was flipped again, and the medium was changed to remove all non-attached cells. After 48h, epithelial cells were seeded on the apical side at 270,000 cells/cm^2^. After 24h, 50/50 medium (half EGM2-MV supplemented and half SAGM supplemented) was used in both monoculture and coculture. Medium was changed daily.

### Measurement of the membrane thickness

The thickness of the membrane was measured with reflective light microscopy. Briefly, the membrane was cut at its centre and imaged using the Axioplan microscope (Zeiss, Oberkochen, Germany). The thickness was measured using Axiovision software. Confocal imaging (z-stack) with LSM710 (Zeiss) was used to confirm the thickness of the membrane. Images were analysed with ImageJ software.

### Transparency

The optical transparency of the membrane was evaluated using a light spectrometer (M1000 Infinite, TECAN, Mannedorf, Switzerland) in the range of 350 to 700nm. Membranes were produced by pipetting a solution of the specific material to be tested on the bottom of a 96-well plate. The volume of the solution was adapted to obtain a 10µm-thin membrane.

### Absorption/adsorption

The ab-and adsorption of small molecules by the membranes was quantified by immerging them in 10µM rhodamine B (Sigma-Aldrich) in PBS for 2h at 37°C. A CE-membrane with ratios 1:1 and 2:1; a collagen membrane; a polyester membrane with 0.4 and 3µm pore sizes; and 40, 10 and 3.5 µm porous PDMS membranes were tested. After immersion in rhodamine B, membranes were washed twice in PBS for 5 min. The fluorescence of each membrane was measured using a standard spectrometer (Infinite M1000, TECAN) with an excitation wavelength of 544 nm and an emission of 576 nm. Pictures of the membranes after immersion were taken with a Leica DMI400 (Leica Microsystems, Buffalo Grove, IL, USA). The PDMS membranes were fabricated by spinning PDMS attached to a silicon wafer at 1650 rpm (resp. 6700 rpm) for 60s to obtain a 40µm (resp. 10µm) membrane. The membrane was then cured for 24h at 60°C. The 3.5µm porous membrane was produced according to a procedure reported previously^9^.

### Permeability assay

Once the cells were confluent, the lower chamber was filled with cell culture medium. The upper chamber was filled with 1μg/mL FITC-Sodium (0.4kDa, Sigma-Aldrich) and 1.5mg/mL RITC-Dextran (70kDa, Sigma-Aldrich) in 50/50 medium (half EGM2-MV supplemented and half SAGM supplemented). The device was incubated for 4h, after which the solution in the upper channel was removed and the top chamber was washed three times with PBS. Subsequently, the solution from the lower chamber was collected. The samples were tested for fluorescence with a multi-well plate reader (M1000 Infinite, TECAN). The FITC-Sodium and RITC-Dextran were excited at 460nm and 553nm, respectively. Emission was measured at 515nm and 627nm, respectively. The permeability of the air-blood barrier was expressed in terms of relative transport, in that the amplitude of the fluorescent signal of the basal chamber solution was normalised to the fluorescence intensity signal of the initial solution of the apical chamber.

### Measurement of deflection

The membrane was cyclically deflected using a homemade electro-pneumatic system generating a cyclic negative pressure that can be tuned from 1 to 30 kPa. The deflection measurement was performed by the evaluation of the height difference between stretched and unstretched membrane. Pressure was applied for 20s, followed by a resting time of 1min. For each membrane, a minimum of four hexagons located at the centre of the membrane were measured. On each hexagon, two points were measured: one at the centre of the membrane and one on the gold mesh hexagon. These values were obtained with an AxioPlan2 Zeiss microscope. Linear stress was calculated based on the absolute deflection of the membrane, which was approximated as a circular segment (Sup. Fig. 12).

### Immunofluorescence

All immunostaining steps were conducted at room temperature. The chips were washed three times with PBS, fixed with 4% paraformaldehyde (Sigma-Aldrich) for 10min and rinsed again three times with PBS. The cells were permeabilised with 0.1% Triton X-100 (Sigma-Aldrich) for 10min and washed three times with PBS. After 45min of blocking in a 2% BSA (Sigma-Aldrich) solution, primary antibodies were diluted in the blocking solution. The chip was incubated for 1.5h. Following incubation, devices were washed three times with PBS, then incubated for 1h with the associated secondary antibody. A 1:2000 dilution of Hoechst was added to image cell nuclei. Finally, the chip was washed with PBS. The top layer was detached from the bottom to image the cells on the membrane. Images were obtained using a confocal microscope (CLSM, Zeiss LSM 710).

### Scanning electron microscope

For SEM acquisition, samples were fixed with 2.5% glutaraldehyde (Merck) in 0.1 M cacodylate buffer (Merck) at pH 7.4 for 1h at room temperature. After rinsing three times in a 0.1M cacodylate buffer, the samples were post-fixed for 10min in a 1% osmium tetroxide solution in 0.1M sodium cacodylate buffer. After rinsing three times with Acqua Dest (Medical Corner 24, Oer-Erkenschwick, Germany), the chips were dehydrated at room temperature in 50%, 70%, 80% and 95% ethanol for 10min each. Next, they were immersed in 100% ethanol three times for 10min. Finally, the samples were immersed in hexamethyldisilane for 10min and then dried at room temperature. Samples were mounted onto stubs with adhesive carbon (Portmann Instruments, Biel-Benken, Germany) and coated by electron beam evaporation with platinum/carbon (thickness of coating: 26nm). Pictures were taken with the DSM982 Gemini digital field emission scanning electron microscope (Zeiss) at an acceleration of 5kV and a working distance of 3 mm.

### Transmission electron microscopy

For TEM acquisition, the chips were fixed with 2.5% glutaraldehyde (Agar Scientific, Essex, UK) in 0.15 M HEPES (Sigma-Aldrich) buffer (670mOsm, pH 7.35). The samples were placed at 4°C. Samples were post-fixed for 1h in a 1% osmium tetroxide solution in 0.1M sodium cacodylate buffer (Merck) and rinsed three times in the same buffer. Next, the chips were dehydrated at room temperature with an ethanol concentration series (70%, 80% and 96%) for 15min each. Then, they were immersed in 100% ethanol (Merck) three times for 10min. The chips were embedded in an epoxy solution and incubated at 60°C for 4 days. For samples without cells, the chips were directly embedded in the epoxy solution. After removing the PDMS surrounding the gold mesh, ultrathin sections (70nm) were cut with an ultramicrotome UC6 (Leica Microsystems) and mounted on 1mm single-slot copper grids. Pictures were taken with a Philips EM 400 transmission electron microscope.

### Numerical simulation

A stationary numerical simulation, using COMSOL Multiphysics 5.3 (COMSOL Multiphysics GmbH, Switzerland), was performed to visualize and illustrate the deformation of the CE-membrane during breathing.

### Statistics

Data are presented as mean ± standard deviation (SD). Two-tailed unpaired Student’s t-test was used to assess the significance of differences. Statistical significance was defined as follows: *p<?0.05, **p<?0.01, ***p<?0.001. Statistical analysis was performed using GraphPad Prism 6 software.

## Acknowledgments

This work was partly supported by the Swiss Innovation Agency (CTI/Innosuisse grant N° 27813.1 PFLS). The authors thank Prof. em. Dr. phil. Nat. Peter Gehr from the Institute of Anatomy of the University of Bern for graciously providing the SEM picture of the lung alveoli (Fig. 1B), as well as Beat Hänni from the same institute for his help processing the samples and taking the TEM pictures (Fig. 5D, 6C & 7B). The authors thank Prof. Dr. med. Vet. Michael Stoffel and Helga Mogel from the Division of Veterinary Anatomy of the University of Bern for their help in the preparation and acquisition of the SEM data (Fig. 5E.F) and the Microscopy Imaging Center (MIC) at the University of Bern. They also thank Jan Schulte for his help in cells surface measurement. Thanks also go to Dr. Usha Sarma for helping revising the manuscript and to Dr. Anne Morbach for the illustration of Fig.1C-G.

## Additional information

Supplementary Information accompanies this paper at “Supplementary Information” file.

## Author contributions

O.T.G. had the original idea of using a hexagonal mesh to mimic the lung alveoli. J.D.S., P.Z., N.H. and O.T.G. designed the experiments. S.A. carried out the proof of concept experiments, P.Z. and S.W. set up the device. P.Z. carried out the experiments and collected the data, except Fig. 3C obtained by SW. P.Z., J.D.S., N.H., O.T.G. analysed and interpreted the data. J.D.S. modelled numerically the deflection of the stretchable membrane. P.Z. and O.T.G. wrote the manuscript. N.S.D., C.M.L. and H.H. provided the study material. All authors reviewed the manuscript.

