## Supplementary information for "Second-generation lung-on-a-chip array with a stretchable biological membrane"

**Supplementary Figure 1: Chip design (made on Solidworks)**
The device was made of a top PDMS layer (apical chamber), a collagen-elastin membrane supported by a gold mesh and a polycarbonate bottom (basolateral chamber). The alignment of the layer was ensured by two dowels pins.

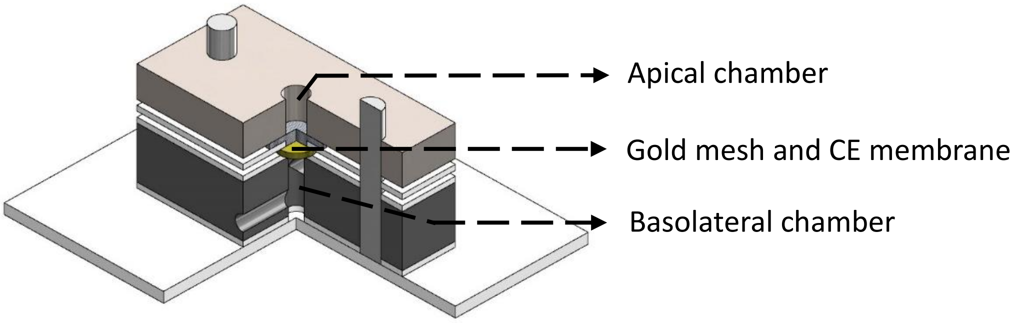


**Supplementary Figure 2: Thickness of the CE-membrane in function of its composition**
The thickness of the CE-membranes made of different ratio of collagen and elastin was evaluated by reflective light. The CE-membrane with a 1:1 (resp. a 2:1) ratio had a final concentration of collagen and of elastin of 3.5mg/mL$^{-1}$ each (resp. 3.5mg/mL$^{-1}$ of collagen and 1.75mg/mL$^{-1}$ of elastin). The collagen membrane was only made of collagen type I with a final concentration of 3.5 mg/mL$^{-1}$. A volume of 1.6µl/mm$^{2}$ was used for each condition. The CE-membrane with a 1:1 ratio had a thickness of 7.7 ± 1.7µm, while that with a 2:1 ratio was 4.9 ± 0.6 µm thin. The collagen membrane was 3.1 ± 0.5µm thin.

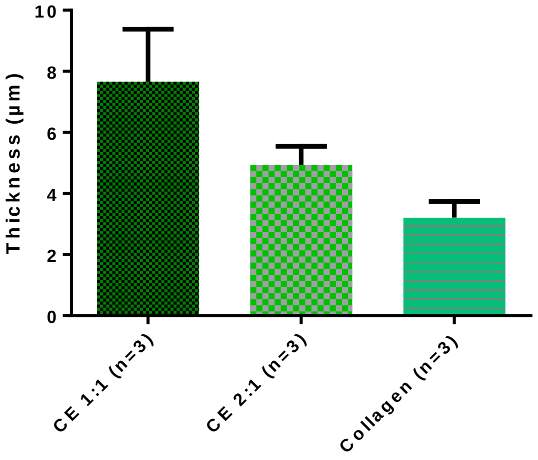

**Supplementary Figure 3: Reproducibility of the CE membranes thicknesses (ratio 1:1, 1.6µL/mm²)**

Several membranes made of collagen and elastin (ratio 1:1) were produced by pipetting 1.6µL/mm² of solution on top of the gold mesh. After drying, their respective thicknesses were evaluated. Four to six measurements were taken in several hexagons across the mesh. The thickness variation across the hexagon array was below 20%. This variation is mainly due to the location of the measurement, as the membrane is thinner in the centre of the hexagon.

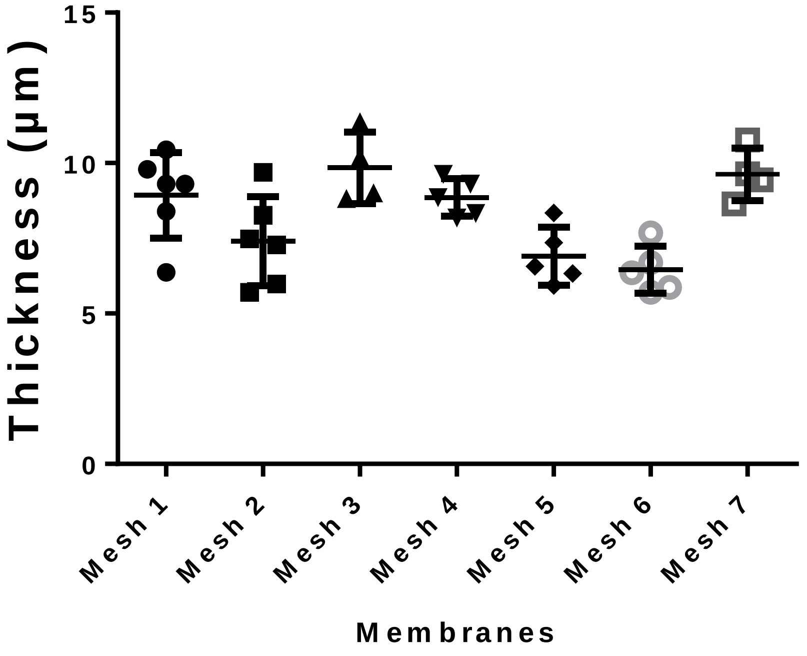


**Supplementary Figure 4: Spectral absorbance of the CE- and of a polyester membrane**

The spectral absorbance of a CE- membrane (10µm-thin with a 1:1 and a 2:1 ratio) and a collagen membrane was smaller than that of a polyester membrane used in standard cell culture inserts.

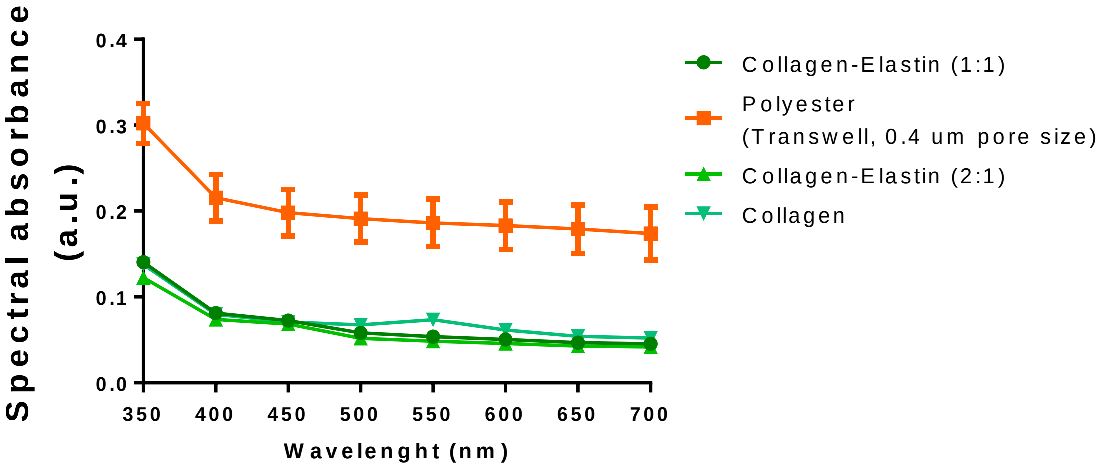

**Supplementary Figure 5: Absorption of Rhodamine B in membranes made of biological and polymeric material**

We evaluated the absorption of Rhodamine B (10µM in PBS) in membranes made of several polymeric and biological materials. The membranes were incubated during 2h in presence of Rhodamine B. It results that the tested polymers (PDMS and polyester) absorbed more Rhodamine B than the tested biological materials. The absorption also varied in function of the thickness of the material and on its porosity. A 40µm-thin PDMS membrane absorbed 2.4 more RhoB molecules than a 10µm-thin PDMS membrane. A Polyester membrane with 3µm pores and a pore density of 2 x $10^6$ pores/$cm^2$, absorbed 2.5 more RhoB molecules than the same membrane with 0.4µm pores and a pore density of 4 x $10^6$ pores/$cm^2$ according to the supplier (https://betastatic.fishersci.com/content/dam/fishersci/en_US/documents/programs/scientific/brochures-and-catalogs/guides/corning-transwell-permeable-supports-guide.pdf).

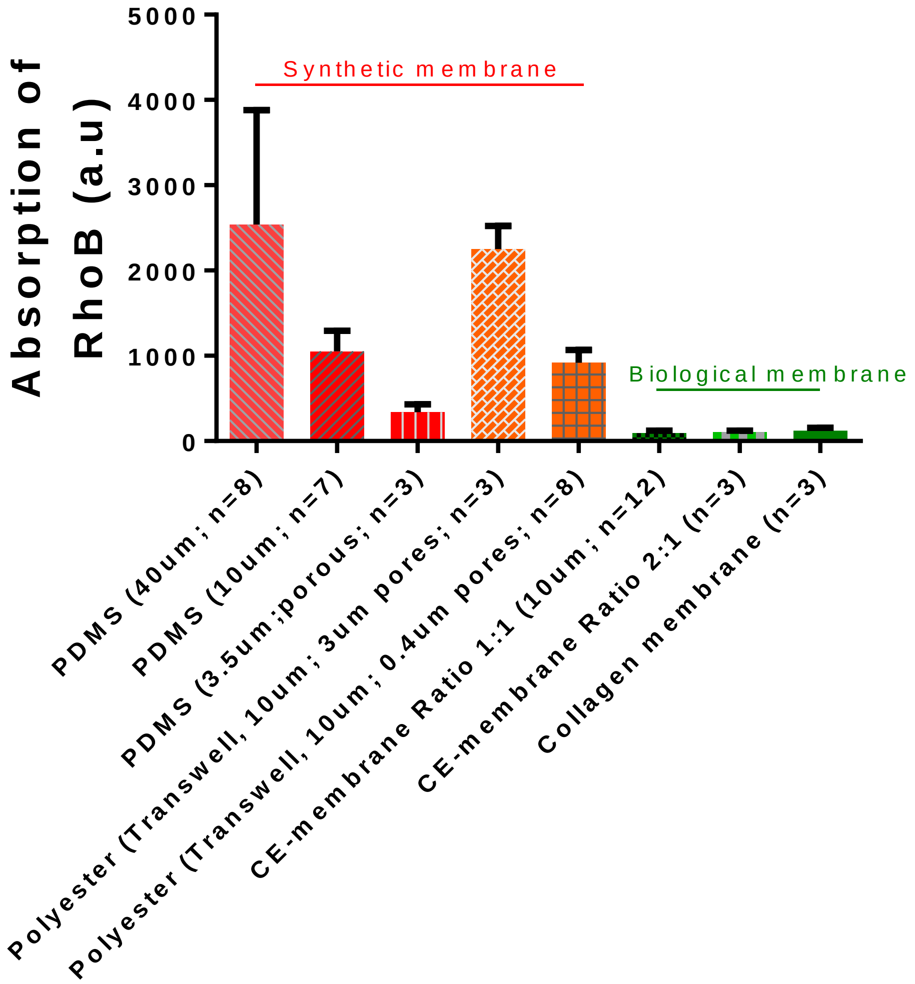

**Supplementary Figure 6: Three-dimensional deflection of the CE-membrane with a monolayer of endothelial cells**

A monolayer of RFP-labelled human lung microvascular endothelial cells, cultured on the CE-membrane (ratio 1:1), was imaged at rest position (no mechanical stress, top view) and stretched with a negative pressure of 2kPa (bottom view).

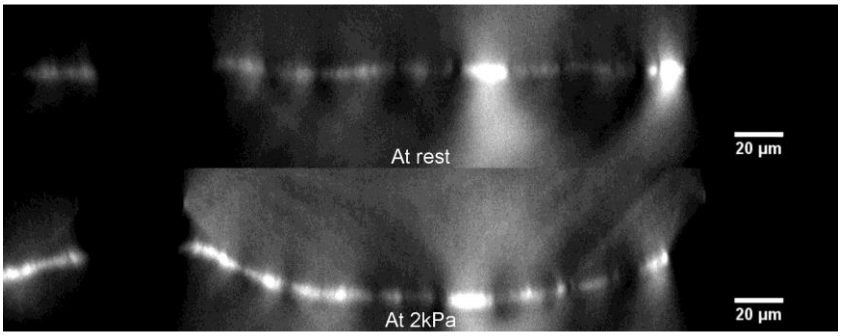


**Supplementary Figure 7: Permeability of the CE-membrane**

We evaluated the transport of FITC-Sodium (0.4kDa) and RITC-Dextran (70kDa) across the CE-membrane (without cells). After four hours of incubation, 25.5 ± 4% of the smaller molecules and 12.0 ± 3.7% of larger molecules were able to cross the membrane and detected in the basolateral chamber of the device.

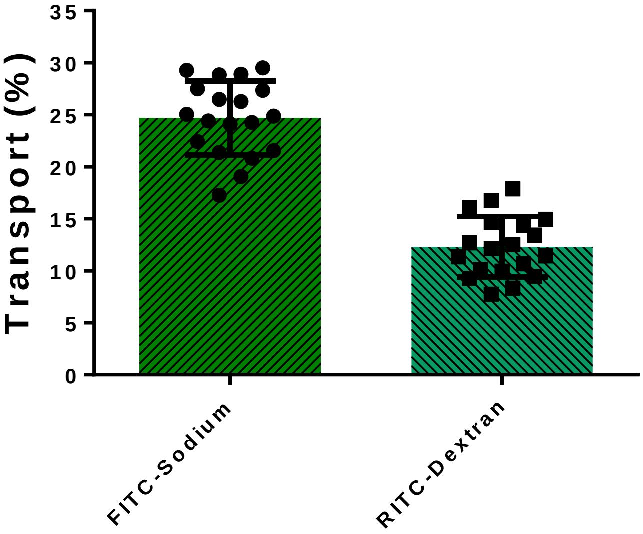

**Supplementary Figure 8: Viability of hAEpC at day 8 after 4 days at ALI**

hAEpC were cultured on the CE membrane in submerged condition (ILI) or Air liquid condition (ALI). In ILI condition, cells were cultured with physiological medium on the apical and the basal chambers for 8 days. In ALI condition, cells were submerged for 4 days, then the medium was removed from the apical side and replaced in the basolateral chamber. The cells were then cultured for 4 days at the air-liquid interface, with the nutrients being provided by diffusion from the basal chamber through the membrane. Culturing the hAEpC at ALI had no impact on the viability of the cells or on the barrier formation, as shown below by the expression of tight junctions (ZO-1).

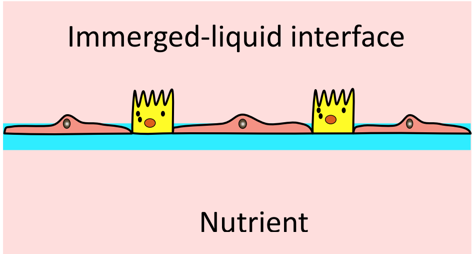


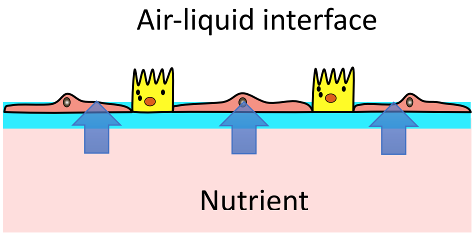


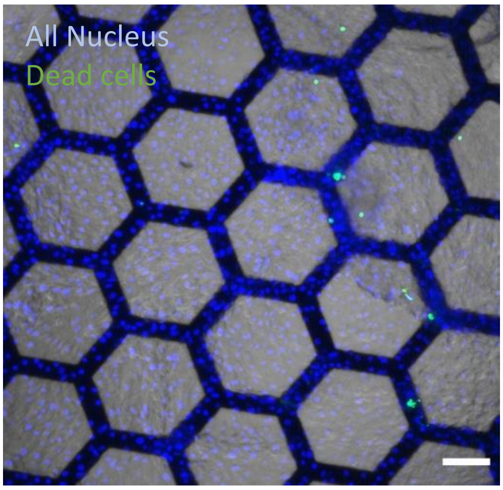


*hAEpC at day 8 in ILI condition; nuclei (blue) and dead cells (green)*

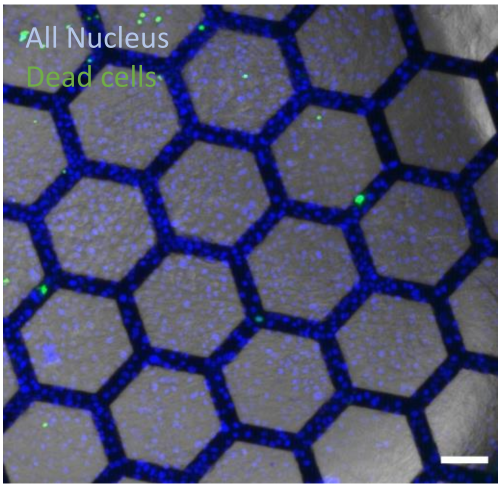


*hAEpC at day 8 in ALI condition; nuclei (blue) and dead cells (green)*

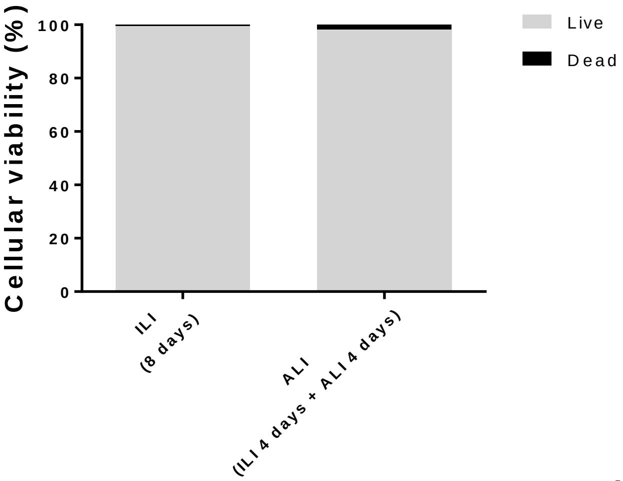


*Viability of hAEpC in monoculture at day 8 in ILI and ALI condition*

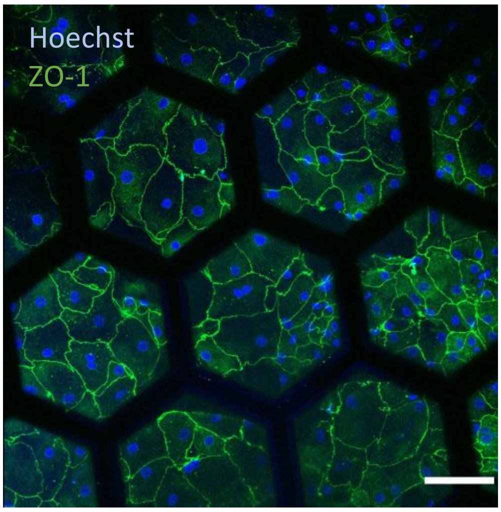


*hAEpC at day 8 in ALI condition; nuclei (Hoechst; blue) and tight junction (ZO-1; green). Scale bar: 100µm*

**Supplementary Figure 9: TEM imaging of hAEpC at different time points**

Human primary alveolar cells (hAEpC) were successfully cultured on the CE-membrane. The cells adherence was excellent on the membrane and did not detach even after one week in culture. Lamellar-like bodies can be seen from day 0 (D0) to day 8 (D8).

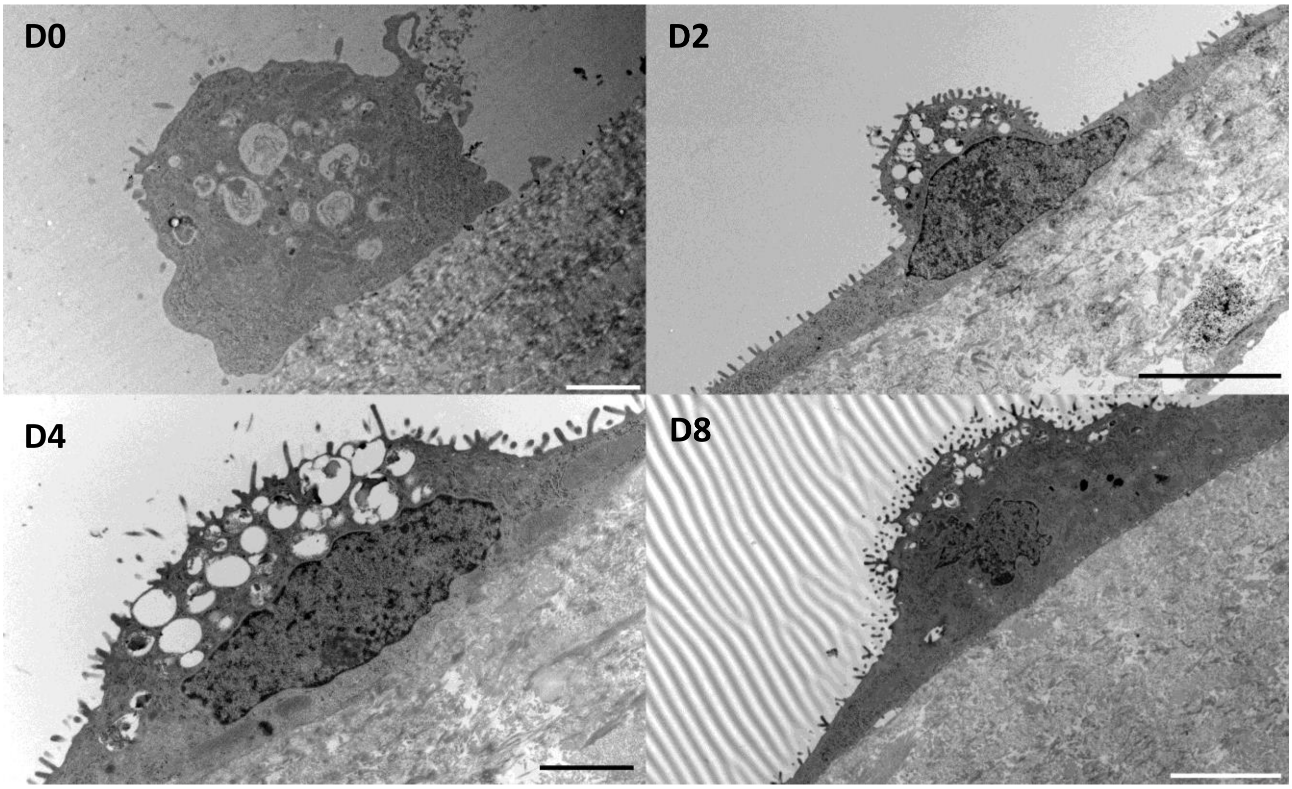


*hAEpC at Day 0. Scale bar: 2µm; Day 2. Scale bar: 5µm; Day 4. Scale bar: 2µm; Day 8. Scale bar: 5µm*

**Supplementary Figure 10: Surface and number of hAEpC in function of the seeding concentration**

The hAEpC were seeded at high concentration (HC, 270'000 cells/cm$^{-2}$) and low concentration (LC, 100'000 cells/cm$^{-2}$). The cellular surface was quantified using ZO-1 (green) to show the cell borders and Hoechst (blue) for the nuclei. When cells were seeded at high concentration, they almost reached confluence at day 2 and their number remains stable (day 8). However, when they were seeded at LC, their number increased significantly between day 2 and day 8.

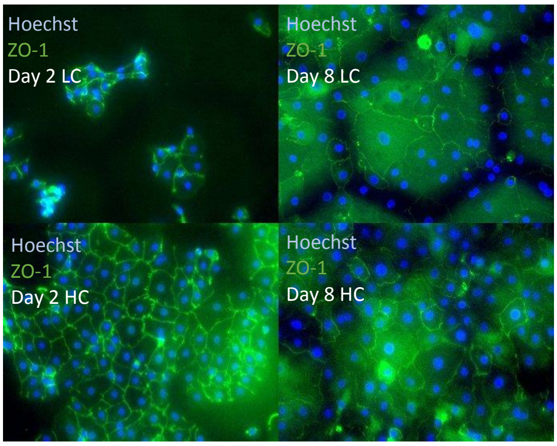


*Day 2 LC (top left); day 8 LC (top right); Day 2 HC (bottom left); Day 8 HC (bottom right)*

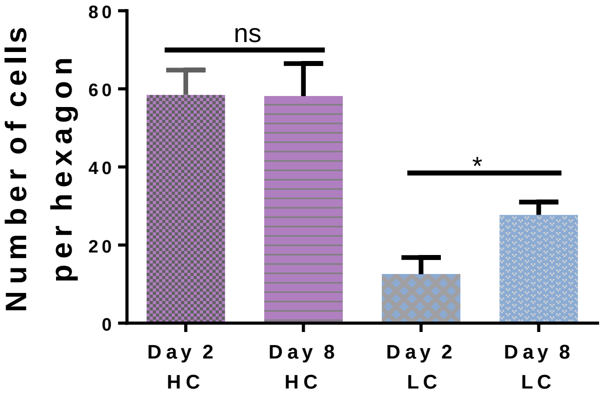

**Supplementary Figure 11: Long-term cultures of hAEpC and of primary endothelial cells**
hAEpC and primary lung endothelial cells were cultured on the CE-membrane for up to 3 weeks without any impact on their viability. At day 21, hAEpC cells still clearly expressed tight junction marker (ZO-1) showing the maintenance of a tight barrier.

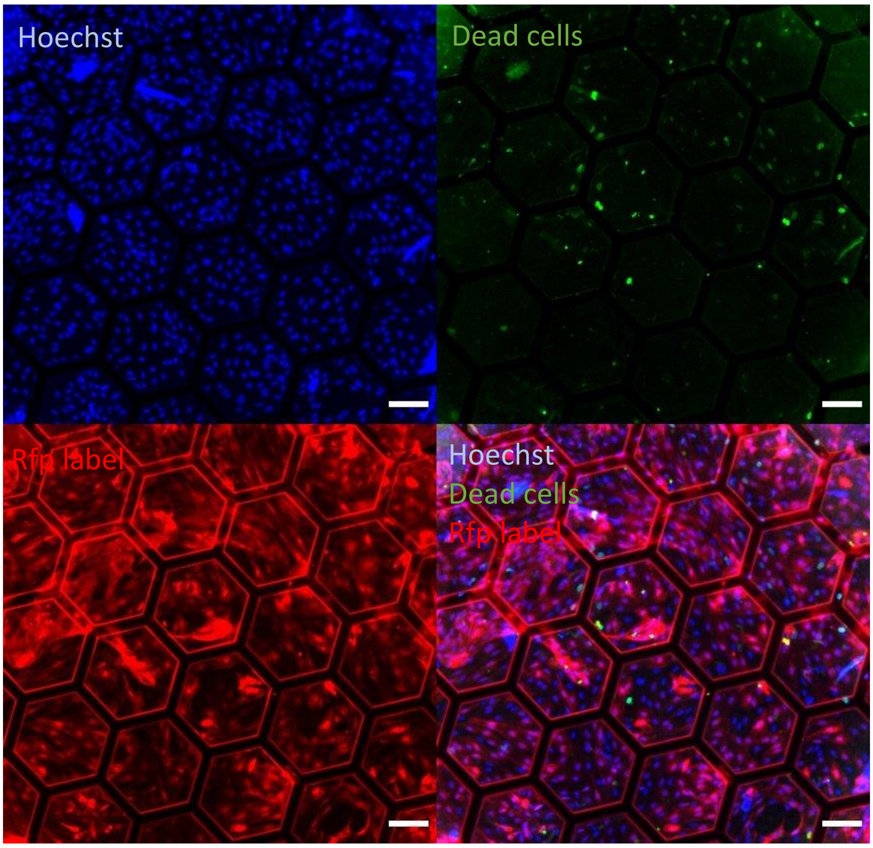


*VeraVec cells at day 21. All nuclei (top left); Dead cells (top right); RFP-label endothelial cells (bottom left); Merged picture (bottom right). Scale bar: 100µm*

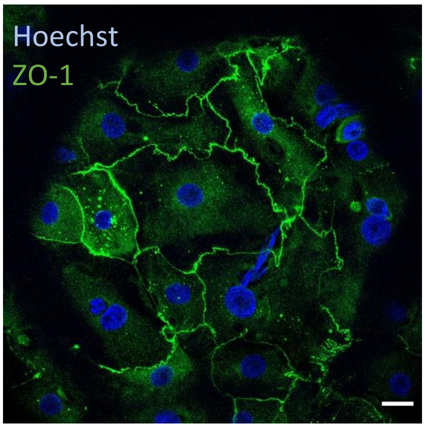


*hAEpC at day 21. ZO-1 (green), Hoechst (Blue). Scale bar: 20µm*

**Supplementary Figure 12: Calculation of the linear mechanical stress**

A simple mathematical model was used to evaluate the mechanical stress of the cells cultured on the membrane. The deflection of the membrane was approximate as a circular segment. On each hexagon, the absolute deflection of the membrane (h) is given by the difference between the deflection of the membrane (1) and the gold mesh (2) (Eq. 1). The length of the stretched membrane (L) can be approximated with a circular segment and it was calculated with the formula (Eq. 2) with (L0) the original length. The change in length (ΔL) can be calculated with the formula (Eq. 3). Finally, the linear strain (ε) is the ratio of change in length to the origin length (Eq. 4).

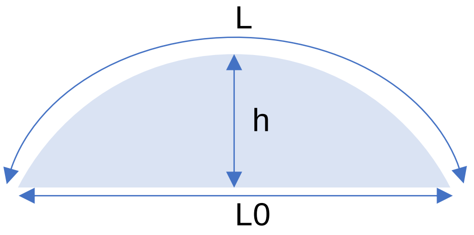


*Equation* 1: $h = h1 - h2$

*Equation* 2: $L = \dfrac{\arctan\left(\frac{2*h}{L0}\right) * (4*h^2 + L0^2)}{2*h}$

*Equation* 3: $\Delta L = L - L0$

*Equation* 4: $\varepsilon\ [\%] = \dfrac{\Delta L}{L0} * 100$

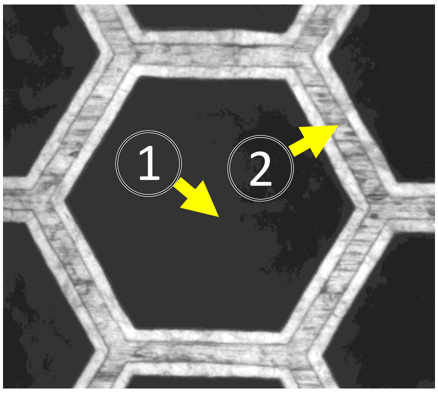


*Measurements points (1: deflection of the CE-membrane; 2: deflection of the gold mesh)*
